# Phase transition of maternal RNAs during vertebrate oocyte-to-embryo transition

**DOI:** 10.1101/2023.05.11.540463

**Authors:** Hyojeong Hwang, Sijie Chen, Meng Ma, Divyanshi, Hao-Chun Fan, Elizabeth Borwick, Elvan Böke, Wenyan Mei, Jing Yang

**Author notes:** Corresponding authors: Jing Yang.

## Abstract

The oocyte-to-embryo transition (OET) is regulated by maternal products stored in the oocyte cytoplasm, independent of transcription. How maternal products are precisely remodeled to dictate the OET remains an open question. In this work, we discover the dynamic phase transition of maternal RNAs during *Xenopus* OET. We have identified 863 maternal transcripts that transition from a soluble state to a detergent-insoluble one after oocyte maturation. These RNAs are enriched in the animal hemisphere and many of them encode key cell cycle regulators. In contrast, 165 transcripts, including nearly all *Xenopus* germline RNAs and some vegetally localized somatic RNAs, undergo an insoluble-to-soluble phase transition. This phenomenon is conserved in zebrafish. Our results demonstrate that the phase transition of germline RNAs influences their susceptibility to RNA degradation machinery and is mediated by the remodeling of germ plasm. This work thus uncovers novel remodeling mechanisms that act on RNAs to regulate vertebrate OET.

## Introduction

The oocyte-to-embryo transition (OET) is one of the most dramatic developmental transitions during which the oocyte and sperm fuse to produce an embryo capable of giving rise to progeny. Prior to the OET, the oocyte accumulates large amounts of maternal products during oogenesis. During the OET, a series of events, including meiotic oocyte maturation, ovulation, fertilization, and zygotic genome activation occur sequentially, allowing the oocyte to transition into a rapidly growing embryo. Strikingly, these events are precisely regulated by maternal gene products stored in fully-grown oocytes, independent of transcription ^1^. After decades of extensive investigation, it is still largely unclear how maternal gene products are remodeled during the OET to direct the initiation of embryonic development.

It is well known that numerous remodeling events happen during the OET. These include the remodeling of cellular organelles, as well as macromolecules in the oocyte. One of the best-studied remodeling events during the OET is the remodeling of the endoplasmic reticulum (ER) during oocyte maturation. In most vertebrate species, the ER is closely associated with the germinal vesicle (GV) with some tubular ER being distributed in the cytoplasm of fully-grown oocytes. After germinal vesicle breakdown (GVBD), the morphology and distribution of the ER are changed massively. As a result, a substantial amount of the ER is placed under the plasma membrane, around the future sperm entry site. This unique organization allows rapid release of calcium from the ER upon sperm entry, facilitating egg activation ^2–10^. In some animal species, ER is pivotal for the asymmetric localization of maternal RNAs, especially some RNAs essential for early embryonic patterning ^11–15^. In *Xenopus*, interfering with ER remodeling during the oocyte maturation severely impairs the proper localization of maternal RNAs ^16^. Interestingly, the proteasome system is remodeled during the OET as well. In mice, proteasomes become highly enriched in the nucleus after the OET ^17^. In *Xenopus*, proteasomes are translocated to the animal hemisphere during the OET. This increases the half-life of vegetally localized Dnd1, allowing the accumulation of Dnd1 protein in the embryo to facilitate germline development ^18^.

A few recent studies demonstrate that at the global level, maternal mRNAs are remodeled during the OET. In humans, many maternal mRNAs are deadenylated first, and then re-polyadenylated. During re-adenylation, massive incorporation of non-A residues occurs. Interfering with this remodeling event arrests human embryos at the 1-cell stage ^19^. In *Xenopus,* a large amount of maternal RNAs is associated with the ER in fully-grown oocytes. During oocyte maturation, the mRNA-ER association is decreased, leading to the relocation of many maternal RNAs in mature eggs. The decreased mRNA-ER association was observed during mouse oocyte maturation as well ^16^. In a study to analyze the dynamic structural changes in the 3’UTR of zebrafish maternal mRNAs, Shi et al. detected an opening in the 3’UTR structure immediately after fertilization and a gradual close during cleavage, followed by a reopening after the zygotic genome activation. These structural changes alter the accessibility to specific RNA-binding proteins, influencing the stability of maternal RNAs ^20^. These findings are exciting and highlight the importance of the remodeling of maternal mRNAs during the OET.

In this study, we report that a subset of maternal mRNA undergoes phase transition during *Xenopus* and zebrafish OET. Our results reveal that some mRNAs encoding cell cycle regulators become more insoluble during the OET, whereas germline RNAs, which are highly insoluble in the oocyte, are solubilized during the OET. We provide evidence that the solubility of germline RNAs is regulated by Xvelo1/Buc proteins, which form the matrix of the germ plasm ^21–24^. Moreover, we show that the solubility of germline RNAs influences their susceptibility to RNA degradation machinery. Our work thus uncovers novel RNA remodeling mechanisms that occur during vertebrate OET.

## Results

### Phase Transition of maternal RNA during oocyte maturation

We recently investigated RNA-ER association during *Xenopus* oocyte maturation by fractionation RNA-seq. Our results reveal that the majority of maternal RNAs are distributed in the cytosolic and ER fractions in the oocyte and mature egg. Interestingly, about 10% of RNAs are present in the insoluble pellet fraction, which is resistant to Triton X-100 extraction and can be precipitated by centrifugation at 800 x g ^16^. While the total amount of RNA in the pellet fraction remains largely unchanged during oocyte maturation, the composition of transcripts in the pellet fraction is dynamically regulated. To simplify our analysis, we combined the cytosolic and ER fractions as the soluble fraction and calculated the percentage of each transcript distributed in the soluble and insoluble (the pellet fraction) fractions. As shown in Fig 1A and S-table 1, while the majority of RNAs are soluble, some RNAs exist in an insoluble form. We detected 233 insoluble transcripts (more than 40% in the pellet fraction) in the oocyte. After oocyte maturation, 29.2% of these transcripts remain insoluble. The remaining 70.8% of transcripts show a decrease in the pellet fraction after oocyte maturation. Among all transcripts mainly distributed in the soluble fraction in the oocyte, 863 transcripts show a significant increase (above 2-fold) in the insoluble fraction after oocyte maturation. These observations indicate that the solubility of maternal RNA is dynamically regulated during oocyte maturation. We categorized RNAs in the insoluble fraction into three groups, RNAs undergoing an **<u>i</u>**nsoluble-to-**<u>s</u>**oluble phase transition (I-S) during oocyte maturation, RNAs going through a **<u>s</u>**oluble-to-**<u>i</u>**nsoluble phase transition (S-I), and RNAs that are highly insoluble in both oocytes and mature eggs (I-I).

**Fig 1.**
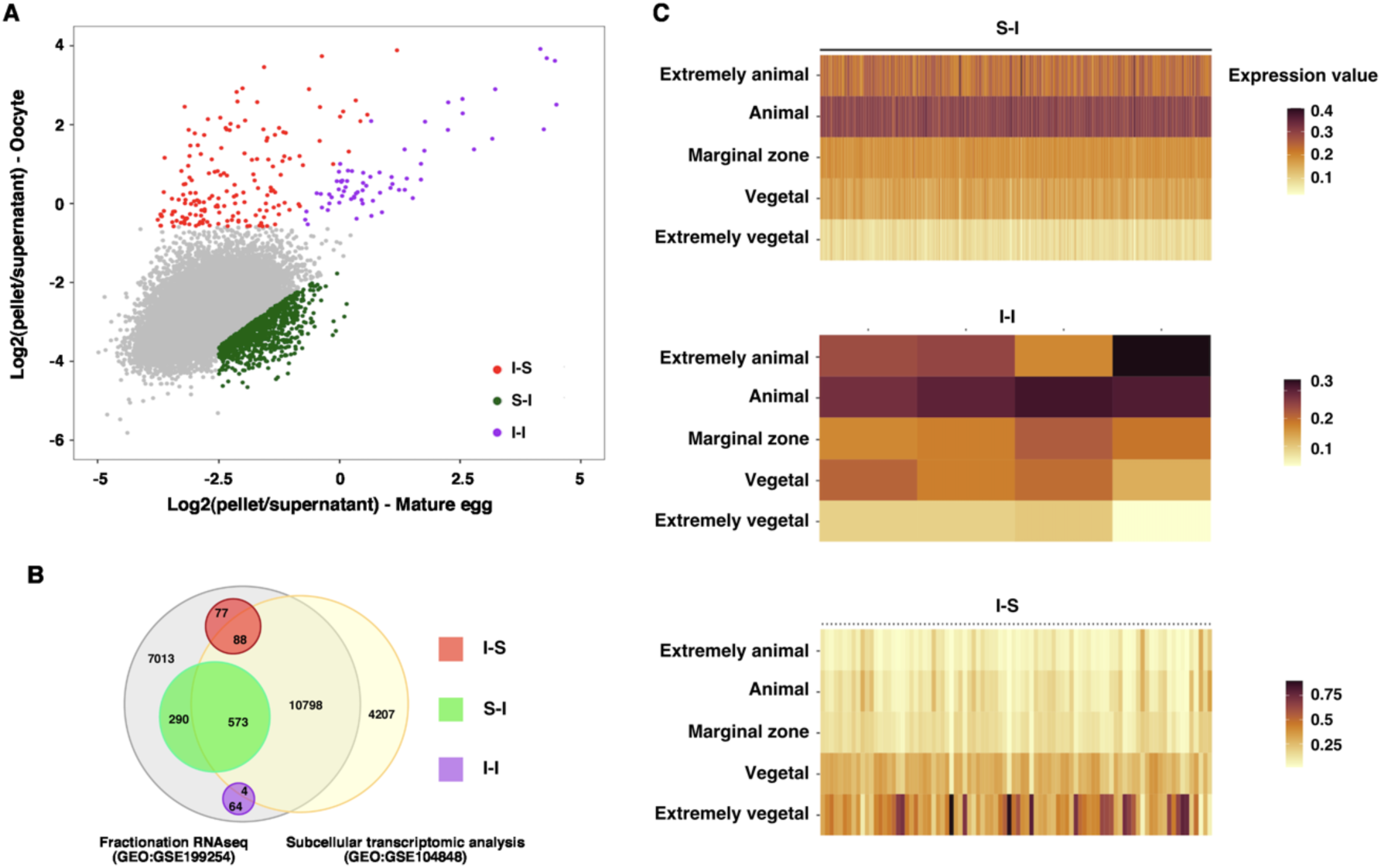
Phase transition of RNA during oocyte maturation. **A.** Scatter plot shows the pellet to the supernatant ratio in the oocyte (Y-axis) and mature egg (X-axis). I-S transcripts (red) were defined as transcripts enriched by more than 40% in the oocyte pellet fraction and reduced by more than 25% in the pellet fraction after oocyte maturation. I-I transcripts (magenta) were defined as those transcripts enriched by more than 40% in the oocyte pellet fraction, excluding those belonging to the I-S group. S-I transcripts (green) were defined as a more than 2-fold increase in the pellet fraction of mature eggs after oocyte maturation and more than 15% pellet enrichment. **B.** Venn diagram shows the overlapping relationship between the fractionation RNA-seq (GEO:GSE199254) and subcellular transcriptomic analysis (GEO:GSE104848). **C**. Heat maps show the distribution of S-I, I-I, and I-S transcripts along the animal-vegetal axis.

In *Xenopus,* many maternal RNAs are asymmetrically distributed along the animal-vegetal axis ^25^. To begin understanding the phase transition of maternal RNAs during oocyte maturation, we first determined if RNAs undergoing phase transitions are asymmetrically located along the A/V axis. To this end, we compared our fractionation RNA-seq data to the subcellular transcriptomic analysis by Sindelka et al. ^26^, in which the distribution of maternal transcripts was analyzed by dissecting the egg along the animal-vegetal axis, followed by RNA-seq. Among 863 S-I RNAs, 573 were detected by Sindelka et al ^26^ (Fig 1B). Intriguingly, all these RNAs are enriched in the animal hemisphere (Fig 1C and S-table 2). In contrast, I-S RNAs show the opposite pattern. Among 88 I-S RNAs detected by Sindelka et al ^26^ (Fig 1B and S-table 2), the majority of them are vegetally localized (Fig 1C and S-table 2). These findings suggest that distinct mechanisms operate along the animal-vegetal axis to regulate the phase transition of maternal RNAs during oocyte maturation.

### Soluble-to-insoluble phase transition of maternal RNA during oocyte maturation

To better understand the RNA phase transition during oocyte maturation, we first studied S-I RNAs. In the fractionation RNA-seq analysis, S-I RNAs show at least a 2-fold increase in the pellet fraction after oocyte maturation (Fig 2A). To validate the RNA-seq result, fully-grown oocytes and mature eggs were fractionated into soluble and insoluble fractions, followed by RT-qPCR for the expression of *ccna1, wee2.S, hmmr.L, parpbp.L, cep152.L, lig4.L, larp1b.S, exd3.L, pif1.L, sox3.S, sema3d.S, ssx2ip.L, espl1.L, wee2.L, ncapd2.S, fbxo43.L, cdc6.L, ccdc18.L, zbtb12.L, kank1.L, ccnb1.2.L, ncbp1.S, eif2ak3.S, dock7.S, ccnb2.L, dbr1.L,* and *rad21.L*. Indeed, all these S-I RNAs show a significant increase in the insoluble fraction after oocyte maturation (Fig 2C and D), confirming these maternal transcripts indeed go through soluble-to-insoluble phase transition during oocyte maturation.

**Fig 2.**
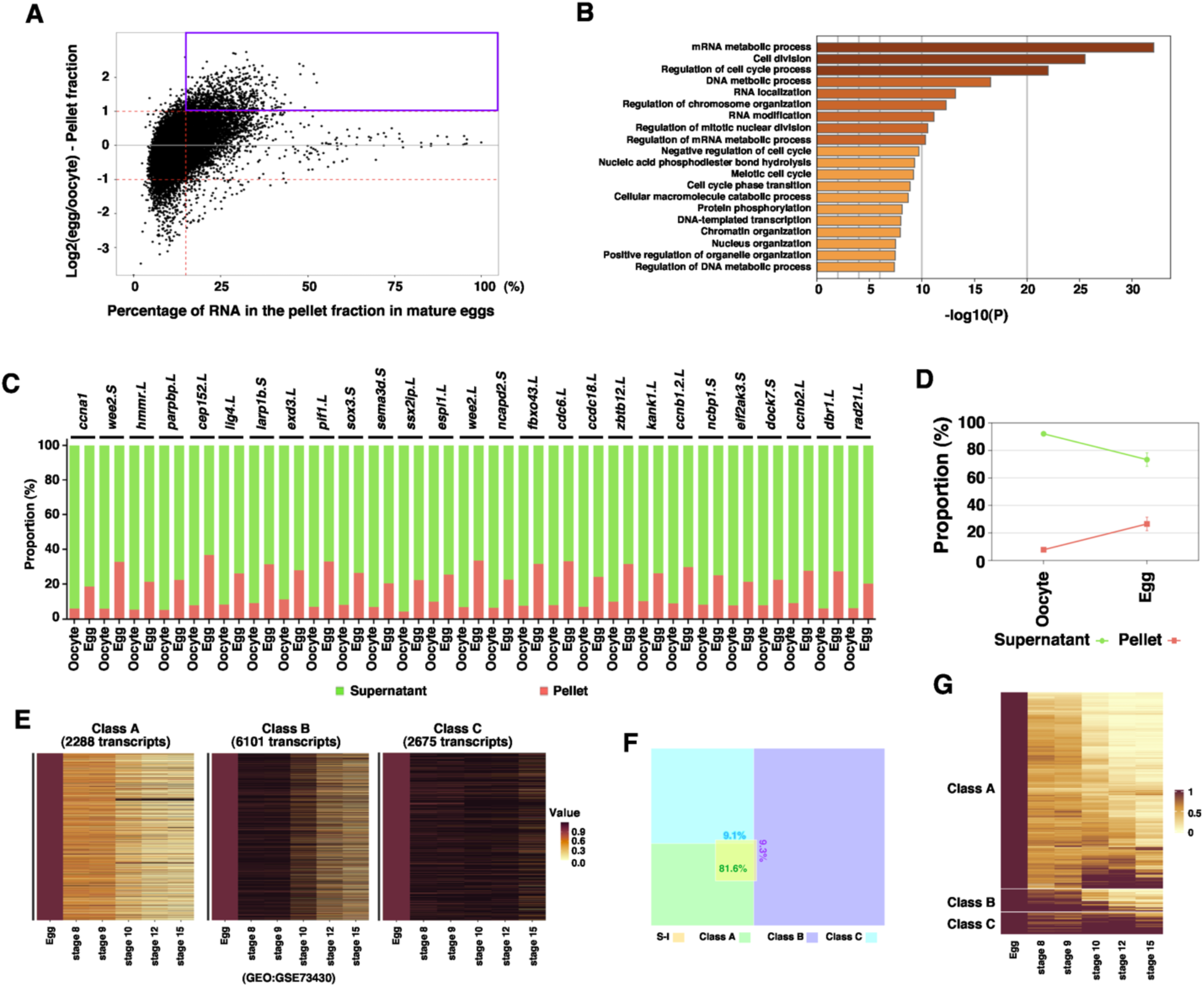
Soluble-to-insoluble phase transition of RNA during oocyte maturation. **A.** MA plot shows the percentage of RNA in the pellet fraction in the mature egg (X-axis) and the ratio between the RNA in the pellet fraction of the egg and that of the oocyte (Y-axis). S-I transcripts are highlighted in the magenta box. **B.** Gene ontology analysis demonstrates the top biological processes that are enriched among S-I transcripts. **C**. Fractionation RT-qPCR was performed to validate fractionation RNA-seq results. The percentage distribution of *ccna1, wee2.S, hmmr.L, parpbp.L, cep152.L, lig4.L, larp1b.S, exd3.L, pif1.L, sox3.S, sema3d.S, ssx2ip.L, espl1.L, wee2.L, ncapd2.S, fbxo43.L, cdc6.L, ccdc18.L, zbtb12.L, kank1.L, ccnb1.2.L, ncbp1.S, eif2ak3.S, dock7.S, ccnb2.L, dbr1.L,* and *rad21.L* in the supernatant and pellet fractions were calculated. **D.** The percentage distribution of all markers analyzed in **C** was combined and plotted into the graph. **E**. Heat maps show the classification of maternal transcripts based on their degradation during the MZT. Class A transcripts are most rapidly degraded. Class B transcripts are degraded relatively slowly. Class C transcripts are relatively stable during early development. **F.** Venn diagram shows the majority of S-I transcripts belong to Class A. **G.** Heatmap shows the expression of S-I RNAs during early embryonic development.

We performed a Gene Ontology (GO) analysis and found that S-I RNAs are involved in the mRNA metabolic process, cell division, DNA metabolic process, and other cellular processes important for cell proliferation (Fig 2B). It is well known that after fertilization, *Xenopus* embryos go through 12 rapid synchronous cell divisions. At the mid-blastula transition (MBT), large-scale transcription happens, followed by slower asynchronous cell divisions ^27, 28^. The lengthening of the cell cycle after the MBT is a consequence of the rapid degradation of maternal cell cycle regulators such as Cyclins during the maternal-to-zygotic transition (MZT) ^29, 30^. Since several cyclin RNAs, including *ccna1*, *ccnb1, ccne1,* and *ccno* undergo soluble-to-insoluble phase transition during oocyte maturation, we set out to determine if the soluble-to-insoluble phase transition correlates with the degradation of maternal RNAs during the MZT. Gene expression profiles of unfertilized *Xenopus* eggs and embryos at various developmental stages have been analyzed by RNA-seq ^31^. Using these datasets, we assessed the turnover of maternal mRNAs during the MZT. We divided maternal RNA into three classes. Class A RNAs are rapidly degraded after the MBT. Degradation of Class B RNAs occurs relatively slowly. Class C RNAs are relatively stable (Fig 2 E, S-table 3). Intriguingly, the majority (81%) of these S-I RNAs fall into Class A (Fig 2F and G), raising the possibility that the soluble-to-insoluble phase transition of RNAs during oocyte maturation may have a correlation with the turnover of these RNAs during the MZT.

### Insoluble-to-soluble phase transition of maternal RNA during oocyte maturation

Next, we investigated RNAs that are enriched in the insoluble fraction in the oocyte. As shown in Fig 3A and S-table 1, 233 I-S transcripts are highly enriched in the pellet fraction in the oocyte. Among these, 165 transcripts undergo an insoluble-to-soluble phase transition during oocyte maturation. We performed GO (Fig 3B) and protein-protein interaction (PPI) analysis (Fig 3C). The results reveal that nearly all *Xenopus* germline RNAs are I-S transcripts. These include *ddx25* ^32^, *nanos1* ^33^, and *dnd1* ^34^, and genes recently found to be important for PGC development ^35^. Many germline regulators form a protein-protein interaction network (Fig 3C).

**Fig 3.**
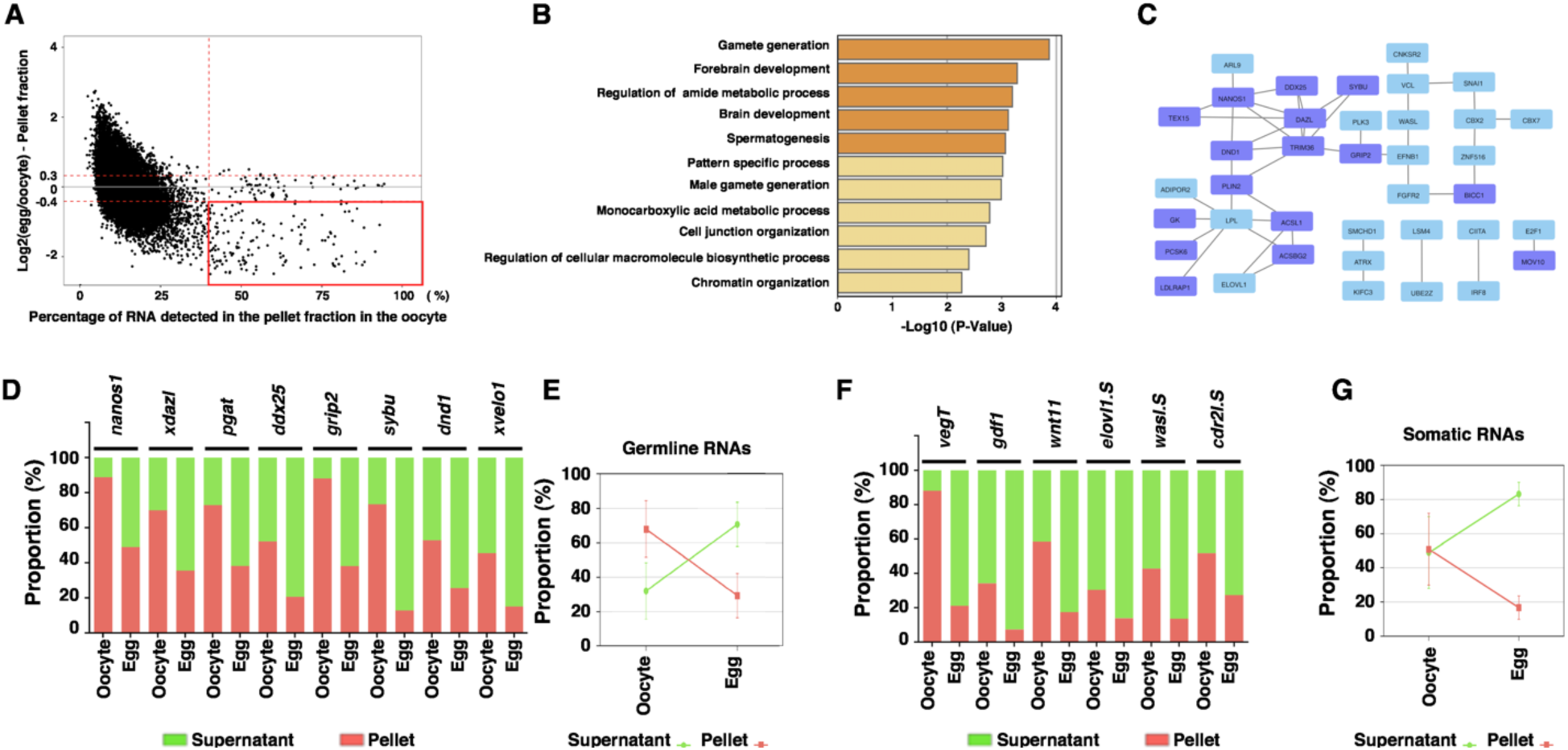
Insoluble-to-soluble phase transition of RNA during oocyte maturation. **A.** MA plot shows the percentage of RNA in the pellet fraction in the oocyte (X-axis) and the ratio between the RNA in the pellet fraction of the egg and that of the oocyte (Y-axis). I-S transcripts are highlighted in the red box. **B.** Gene ontology analysis demonstrates the top biological processes that are enriched among I-S transcripts. **C.** Protein-Protein Interaction (PPI) map shows the transcripts selected from a red box in panel **A**. This PPI shows only proteins interacting with at least one or more other proteins. Purple boxes indicate germline transcripts. **D-G**. Fractionation RT-qPCR was performed to validate fractionation RNA-seq results. **D.** The percentage distribution of germline I-S RNAs, including *nanos1, xdazl, pgat, ddx25, grip2, sybu, dnd1,* and *xvelo1* in the supernatant and pellet fractions, were calculated. **E.** The percentage distribution of all germline RNAs analyzed in **D** was combined and plotted into the graph. **F.** The percentage distribution of somatic I-S RNAs, including *vegT, gdf1, wnt11, elov11.S, wasl.S,* and *cdr2l.S* in the supernatant and pellet fractions, were calculated. **G.** The percentage distribution of all somatic I-S RNAs analyzed in **F** was combined and plotted into the graph.

To validate RNA-seq data experimentally, we fractionated *Xenopus* oocytes and mature eggs into the soluble and insoluble fractions and analyzed 8 germline I-S RNAs (*nanos1*^33^*, xdazl*^36^*, pgat*^37^, *ddx25*^32^*, grip2*^38^*, sybu*^39^*, dnd1*^34^, and *xvelo1*^40^) and 6 somatic I-S RNAs (*vegT, gdf1, wnt11, elovl1.S, wasl.S*, and *cdr2l.S*) by RT-qPCR. Indeed, all the germline transcripts analyzed are enriched in the insoluble fraction in the oocyte. Among these, *nanos1* and *grip2* RNAs represent the most extreme cases, with as high as 80% of RNAs distributed in the insoluble fraction (Fig 3D and E). After oocyte maturation, the percentage of *nanos1*, *xdazl*, *pgat*, *ddx25*, *grip2*, *sybu*, *dnd1*, and *xvelo1* mRNAs in the insoluble fraction decreases more than two-fold (Fig 3D and E). Similar to germline I-S RNAs, *vegT, gdf1, wnt11, elovl1.S, wasl.S*, and *cdr2l.S* are enriched in the insoluble fraction in the oocyte, and are released into the soluble fraction after oocyte maturation (Fig 3F and G). In parallel, we assessed the expression of I-I RNAs by fractionation RT-qPCR. We found *rgs2.L, rnu2, thbs1.S, gata6*, and *bcam.S* are enriched in the insoluble fraction in both oocyte and mature egg (S-Fig 1).

We further extended our analysis by determining when germline mRNAs become insolubilized during oogenesis, we collected stage II, III, IV, and VI oocytes, and performed fractionation RT-qPCR. Our results reveal that *nanos1, pgat, xdazl, ddx25, sybu*, and *grip2* are significantly enriched in the insoluble phase as early as in stage II oocytes (S-Fig 2A). In contrast, *dnd1* and *xvelo1* are initially soluble and gradually move into the insoluble fraction as oogenesis proceeds (S-Fig 2A). The timing of germline RNAs insolubilization correlates with the assembly of these RNAs into the Balbiani body (Bb) or germ plasm (S-Fig 2B). As a control, we also assessed I-I RNAs. Our results reveal that *thbs1.S, gata6*, and *rgs2.L* are highly enriched in the insoluble fraction throughout the oogenesis. Interestingly, the majority of *rnu2* transcript was detected in the insoluble fraction during early oogenesis. In stage IV and VI oocytes, the percentage of *rnu2* in the insoluble fraction decreases. By contrast, *bcam.S* is soluble during early oogenesis, but becomes increasingly insoluble in stage IV and VI oocytes (S-Fig 2C). These results suggest that different RNAs are recruited into some insoluble compartments in the oocyte through distinct mechanisms during oogenesis.

### Insoluble-to-soluble phase transition of germline RNAs is a consequence of the degradation of Xvelo1

To understand the biological significance of RNA phase transition during oocyte maturation, we investigated the mechanism governing the insoluble-to-soluble phase transition of germline RNAs. As germline RNAs are sequestered in the germ plasm in the oocyte, we hypothesized that the germ plasm is remodeled during oocyte maturation, leading to the phase transition of germline RNAs.

Since Xvelo1 and its zebrafish homolog Buc are essential for the formation of Balbiani body (Bb) and germ plasm, we examined the expression of Xvelo1 during oocyte maturation. Our results reveal that the Xvelo1 protein is highly enriched in the insoluble fraction in the oocyte. Even after immunoprecipitation (IP) enrichment, we still could not detect Xvelo1 protein in the soluble fraction (Fig 4A). After oocyte maturation, the expression level of Xvelo1 is decreased sharply. After fertilization, the level of Xvelo1 is further reduced. We could no longer detect the Xvelo1 protein after the mid-blastula transition (MBT) (Fig 4B). To confirm this finding, we performed immunostaining. In fully grown oocytes, we were able to detect numerous Xvelo1 puncta in the cortical region and deep cytoplasm in the vegetal hemisphere. After germinal vesicle breakdown (GVBD), the number of Xvelo1 puncta is reduced gradually. In mature eggs, we could detect Xvelo1 puncta only in the cortex of the vegetal pole (Fig 4C).

**Fig 4.**
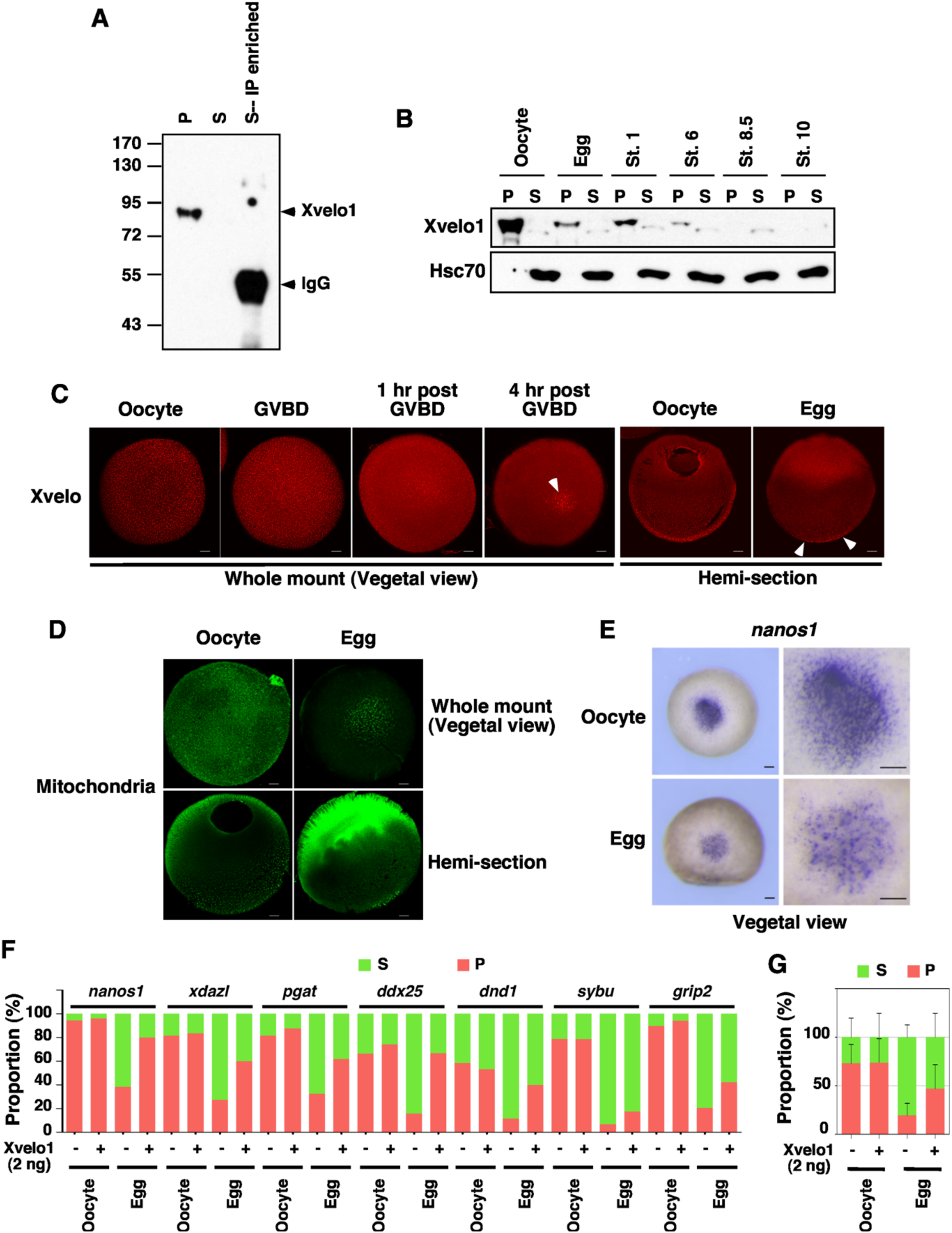
Turnover of Xvelo1 during oocyte maturation results in the solubilization of germline RNAs. **A.** Xvelo1 protein is enriched in the insoluble fraction in the oocyte. Oocytes were lysed in NP-40 lysis buffer. After centrifugation, the lysate was separated into the supernatant (S) and pellet (P). Supernatant prepared from 10 oocytes was incubated with an anti-Xvelo1 antibody to enrich Xvelo1 in the soluble fraction. The supernatant, pellet, and IP samples were mixed with SDS sample buffer and subjected to western blotting. **B.** The expression of Xvelo1 in the oocytes, ovulated eggs, and embryos at stages 1, 6, 8.5, and 10 was analyzed by western blot. **C.** The expression of Xvelo1 during oocyte maturation was analyzed by IF. White arrowheads point to Xvelo1 remaining in the vegetal pole in mature eggs. The scale bars indicate 100 µm. **D**. Oocytes and mature eggs of the Dria transgenic frogs, which carry a mitochondria-specific GFP transgene, were stained with an anti-GFP antibody. The scale bars indicate 100 µm. **E**. Whole mount *in situ* results show *nanos1* transcripts are located in punctate aggregates in the vegetal of the oocyte. After oocyte maturation, *nanos1* transcripts show a diffuse appearance, with only a small number of puncta remaining in the vegetal pole. The scale bars indicate 200 µm. **F** and **G**. Overexpression of Xvelo1 prevents solubilization of germline RNAs after oocyte maturation. Oocytes were injected with 2 ng of *Xvelo1* RNA, and cultured for 2 days, followed by progesterone treatment. Fractionation RT-qPCR was performed to assess the phase transition of *nanos1, xdazl, pgat, ddx25, dnd1, sybu*, and *grip2* (**F**). **G** is the combination of all germline RNAs analyzed in **F**.

As Xvelo1 is the key component of the germ plasm matrix ^21–24^, we went on to determine if the germ plasm is indeed remodeled during oocyte maturation. We assessed mitochondria and *nanos1* RNA, which are sequestered in the germ plasm in the oocyte. Consistent with the dynamic changes in Xvelo1 expression, we detected a large amount of mitochondria aggregates in the entire vegetal hemisphere. After oocyte maturation, mitochondria aggregates were detected only in the cortex at the vegetal pole (Fig 4D). Similarly, *nanos1* RNAs form puncta in the vegetal pole of fully-grown oocytes. After oocyte maturation, the number of *nanos1* puncta is reduced. Diffused *nanos1 in situ* signals become obvious in the vegetal pole of mature eggs (Fig 4E).

The above results support the idea that degradation of Xvelo1 during oocyte maturation results in the insoluble-to-soluble phase transition of germline RNAs. To test this hypothesis directly, we asked if overexpression of Xvelo1 could prevent the solubilization of germline RNAs during oocyte maturation. Indeed, we found overexpression of Xvelo1 significantly decreased the solubility of germline RNAs in mature eggs (Fig 4F and G). Taken together, we conclude that the phase transition of germline RNAs is a consequence of the degradation of Xvelo1 during oocyte maturation.

### Insoluble germline RNAs are resistant to RNase A treatment *in vitro*

In *Xenopus* oocytes, germline determinants are sequestered in numerous small germ plasm “islands” that are finely dispersed in the vegetal hemisphere. After fertilization, they coalesce into a few large aggregates that are inherited later by primordial germ cells (PGCs). Germline components remaining in the somatic tissue are degraded during the maternal-to-zygotic transition. When overexpressed in early zebrafish embryos, Buc, which is the homolog of Xvelo1, forms aggregates and prevents clearance of germline RNAs in somatic tissue, converting somatic cells into PGCs ^22, 41^. We speculated that germline RNAs sequestered in the germ plasm may be protected from RNA degradation machinery. Solubilization of germline RNAs during oocyte maturation increases their chance for degradation, facilitating clearance of germline RNAs in the soma during embryonic development.

To test the above hypothesis, we carried out an *in vitro* RNase A treatment assay. We crushed oocytes in lysis buffer and treated the crude lysates with various amounts of RNase A at 37 C° for 5 min. RT-qPCR was performed subsequently to measure the sensitivity of *nanos1, ddx25, grip2, actin, hsc70*, and *h2a* RNAs to RNase A. Indeed, we found that compared to somatic RNAs (*actin, hsc70*, and *h2a*), germline RNAs (*nanos1, ddx25,* and *grip2)*, which are highly insoluble in the oocyte, are less sensitive to RNase A treatment (Fig 5A). We carefully assessed the changes in the sensitivity of RNAs to RNase A during oocyte maturation. Crude lysates from oocytes and mature eggs were treated with 12.5 pg/µl of RNase A for various amounts of time, followed by RT-qPCR for a larger panel of germline RNAs (*nanos1, xdazl, pgat, sybu, dnd1, xvelo1, ddx25,* and *grip2*) and somatic RNAs (*actin, psma1, odc, gapdh, psme3, ccna1, hsc70*, and *h2a*) (Fig 5B). As expected, somatic mRNAs in oocytes and mature eggs were rapidly degraded by RNase A. The sensitivity of somatic RNAs to RNase A remains unchanged during oocyte maturation. Compared to somatic mRNAs, germline mRNAs were degraded more slowly and to a lesser extent. Interestingly, we found that germline RNAs became more sensitive to the RNase A treatment after oocyte maturation (Fig 5C). To determine if the change in the sensitivity of germline RNAs to RNase A is a result of the solubilization of germline RNAs during oocyte maturation, we fractionated RNase A-treated lysates and measured the sensitivity of soluble and insoluble germline RNAs to RNase A (Fig 5D). As shown in Fig 5E, soluble germline RNAs are highly sensitive to RNase A treatment. We could not detect any difference between oocytes and mature eggs. Insoluble germline RNAs are more resistant to RNase A treatment. Among 8 germline RNAs analyzed, the sensitivity of insoluble *nanos1, xdazl, pgat, sybu, xvelo1,* and *ddx25* to RNase A remains unchanged after oocyte maturation. The sensitivity of insoluble *grip2* and *dnd1* to RNase A increases slightly. Collectively, the above results suggest that when sequestered in the germ plasm, germline RNAs are less prone to degradation. Solubilization of germline RNAs increases their susceptibility to RNA degradation machinery.

**Fig 5.**
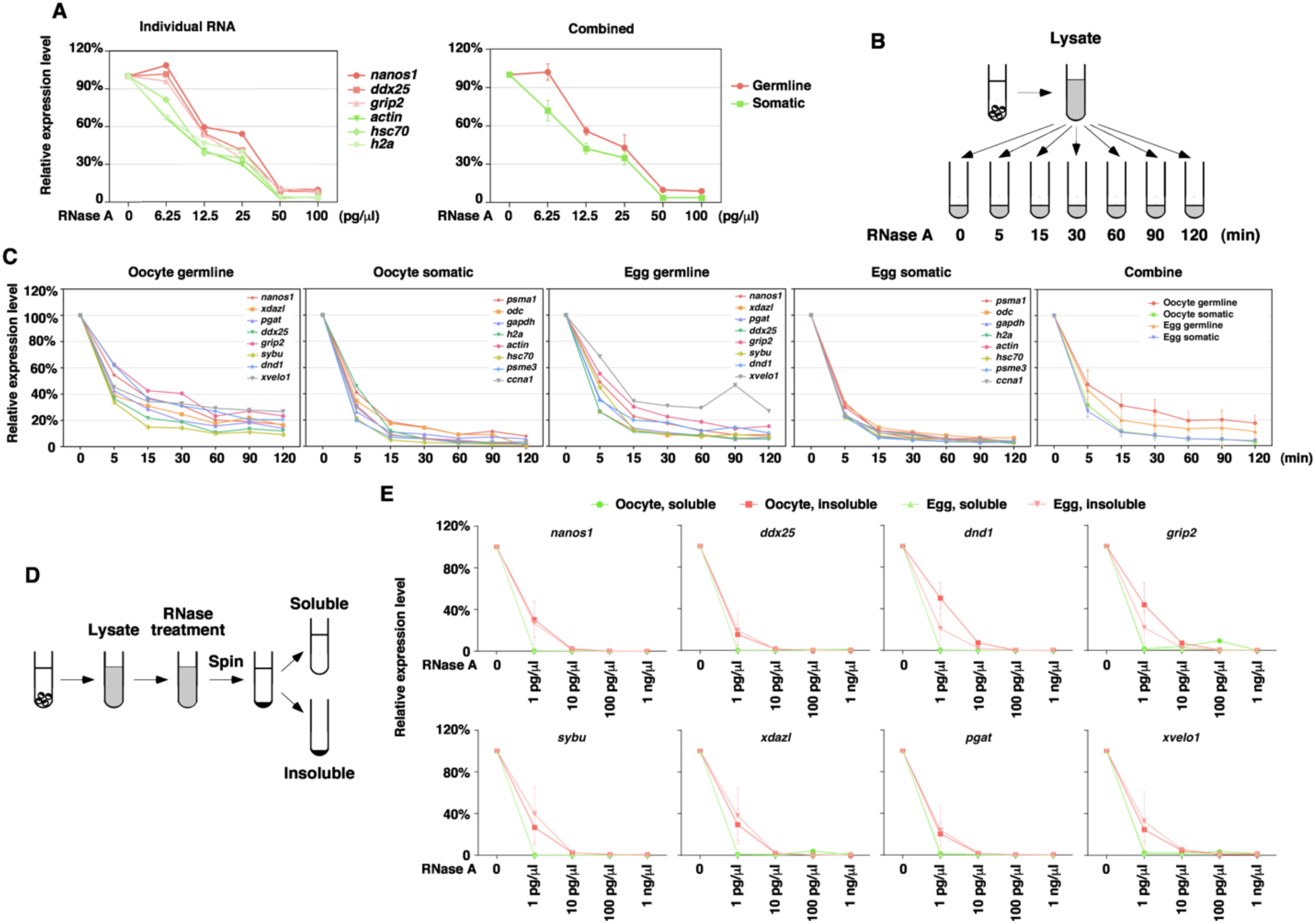
Insoluble germline RNAs are resistant to RNase A *in vitro*. **A.** Crude oocyte lysate was treated with various doses of RNase A. After RNase A-treatment, RNA was extracted for RT-qPCR. The level of *nanos1, ddx25, grip2, actin, hsc70*, and *h2a* was measured. RNA from untreated lysate was set as 100%. **B.** Schematic diagram shows the procedure for the experiments in panel **C**. **C.** Crude oocyte lysate was treated with 12.5 pg/μl RNase A for various amounts of time. Degradation kinetics of germline (*nanos1, xdazl, pgat, ddx25, grip2, sybu, dnd1*, and *xvelo1*) and somatic (*psma1, odc, gapdh, h2a, actin, hsc70, psme3,* and *ccna1*) RNAs were measured by RT-qPCR. The expression of each germline and somatic RNAs in the oocyte and the mature egg was shown individually. The panel on the right side is the combination of all germline and somatic RNAs in the oocyte and egg. **D.** Schematic diagram shows the procedure of experiments in panel **E**. **E.** Crude lysate was treated with RNase A, separated into the soluble and insoluble fractions, followed by RT-qPCR for *nanos1, xdazl, pgat, ddx25, grip2, sybu, dnd1*, and *xvelo1*.

### Solubility and stability of germline RNAs in zebrafish *bucky ball* mutant

To definitively determine if the germ plasm, by maintaining germline RNAs in an insoluble state, protects them from the RNA degradation machinery, we took advantage of the zebrafish *bucky ball* mutant, which lacks Buc protein and is deficient in the formation of Bb and germ plasm. As reported previously ^21, 22, 42^, the ER and mitochondria are accumulated in the Bb in wild-type zebrafish oocytes, but are randomly distributed in the entire cytoplasm in *bucky ball* mutant oocytes (Fig 6A). We performed fractionation and assessed the solubility of several germline RNAs, including *nanos3* ^43^*, dnd* ^44^*, dazl* ^45^*, ca15b* ^46^*, gra* ^47^, and *vasa* ^48^. We found these germline RNAs are enriched in the insoluble fraction in fully-grown wild-type zebrafish oocytes. In contrast, the amount of germline RNAs detected in the insoluble fraction is markedly reduced in *bucky ball* mutant oocytes (Fig 6B), demonstrating that Buc plays an important role in maintaining zebrafish germline RNAs in an insoluble state. We further compared the expression of germline RNAs in control and *bucky ball* mutant embryos at the 1-cell stage. We detected a statistically significant decrease in the expression of *vasa, gra*, and *dazl* (Fig 6C). Since *bucky ball* mutant embryos cannot form blastoderm properly, we could not analyze the degradation of germline RNAs during the MZT. Nevertheless, the observation that the expression of a subset of germline RNAs is decreased in *bucky ball* mutant embryos at the 1-cell stage supports the idea that soluble germline RNAs are more susceptible to RNA degradation machinery.

**Fig 6.**
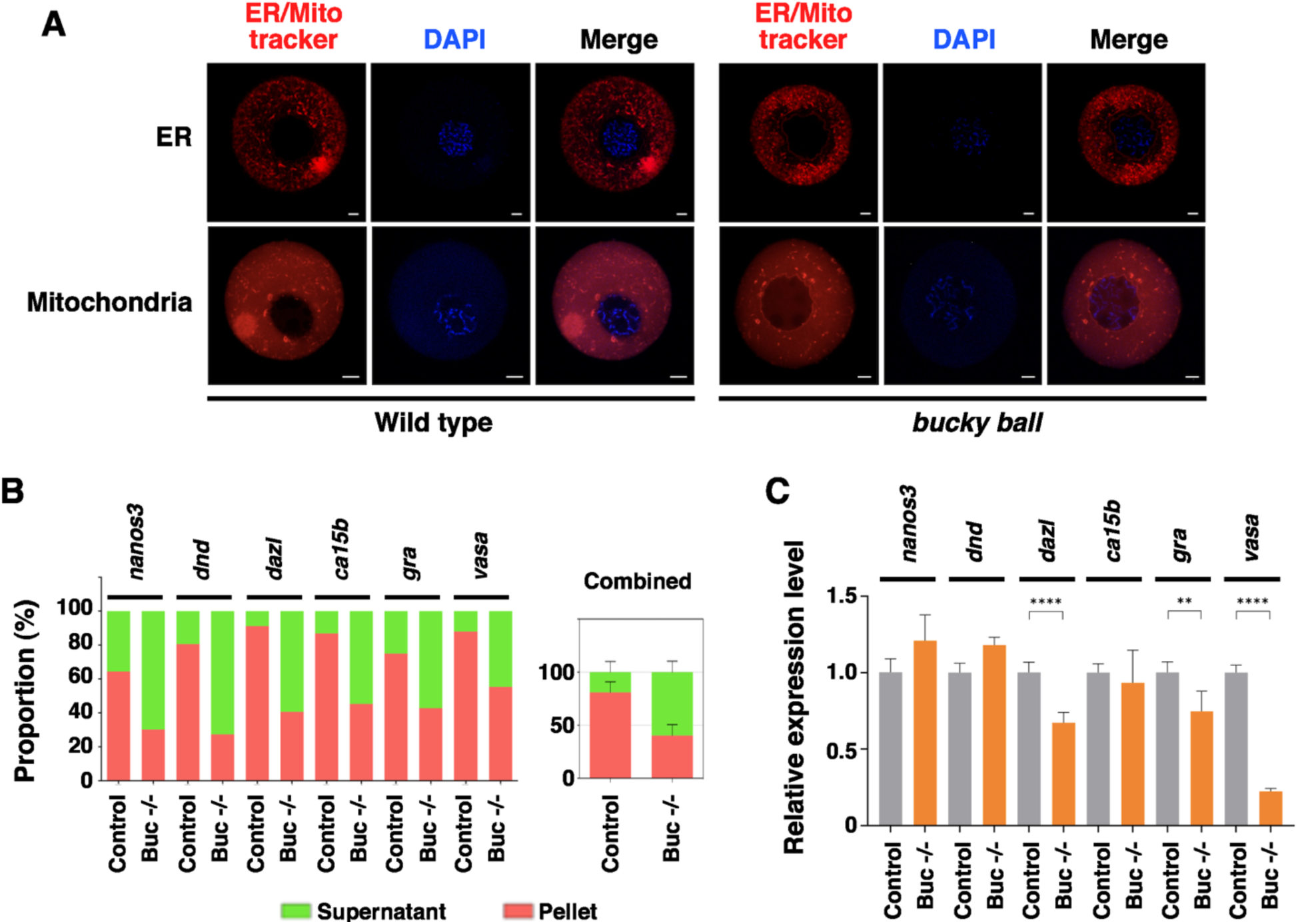
Buc regulates the solubility and stability of germline RNAs in zebrafish. **A.** Wild type and *bucky ball* oocytes were stained with ER-tracker and Mito-tracker. Scale bars on the images for ER trackers indicate 20 µm, and those for Mito trackers indicate 10 µm. **B.** Fractionation RT-qPCR was performed to measure the solubility of *nanos3, dnd, dazl, ca15b, gra*, and *vasa* in fully-grown oocytes from the wild-type fish and *bucky ball* mutants. The right panel is the combination of all these germline RNAs. **C.** The expression of *nanos3, dnd, dazl, ca15b, gra*, and *vasa* in 1-cell stage embryos derived from the wild-type and *bucky ball* mutant females were assessed by RT-qPCR. Student’s *t*-tests were performed. ** p<0.01, ****p<0.0001.

Inspired by the above findings, we went on to investigate if the phase transition of mRNA occurs during zebrafish OET. We performed fractionation on oocytes at various stages, ovulated eggs, and pre-MBT embryos, and assessed the solubility of several germline RNAs, together with *ccna1, ccnb1*, and *wee2,* which are S-I RNAs in *Xenopus* (Fig 2). Our results reveal that the solubility of *ccna1* and *ccnb1* decreases after oocyte maturation. In the case of *wee2*, although its solubility is not significantly changed during oocyte maturation, it is more enriched in the insoluble fraction after fertilization (Fig 7). Germline RNAs show a rather complex pattern. Compared to somatic RNAs, germline RNAs are more enriched in the insoluble fraction in fully-grown oocytes. Among the six germline RNAs analyzed, *dazl, ca15b, gra*, and *vasa* are already in the insoluble fraction in stage I oocytes. Recruitment of *nanos3* and *dnd* to the insoluble fraction occurs gradually during oogenesis. We did not detect any significant changes in the solubility of germline RNAs during oocyte maturation. Instead, we found that the solubility of *dazl, ca15b, nanos3*, and *dnd* increases gradually during pre-MBT stages (Fig 7). Thus, similar to what happens in *Xenopus*, some cell cycle regulators become increasingly insoluble during zebrafish oocyte maturation and early development. The insoluble-to-soluble phase transition does occur in some zebrafish germline RNAs, albeit during cleavage and blastula stages. While some species-specific differences clearly exist, the above results demonstrate that the phase transition of maternal mRNA during the OET is evolutionarily conserved.

**Fig 7.**
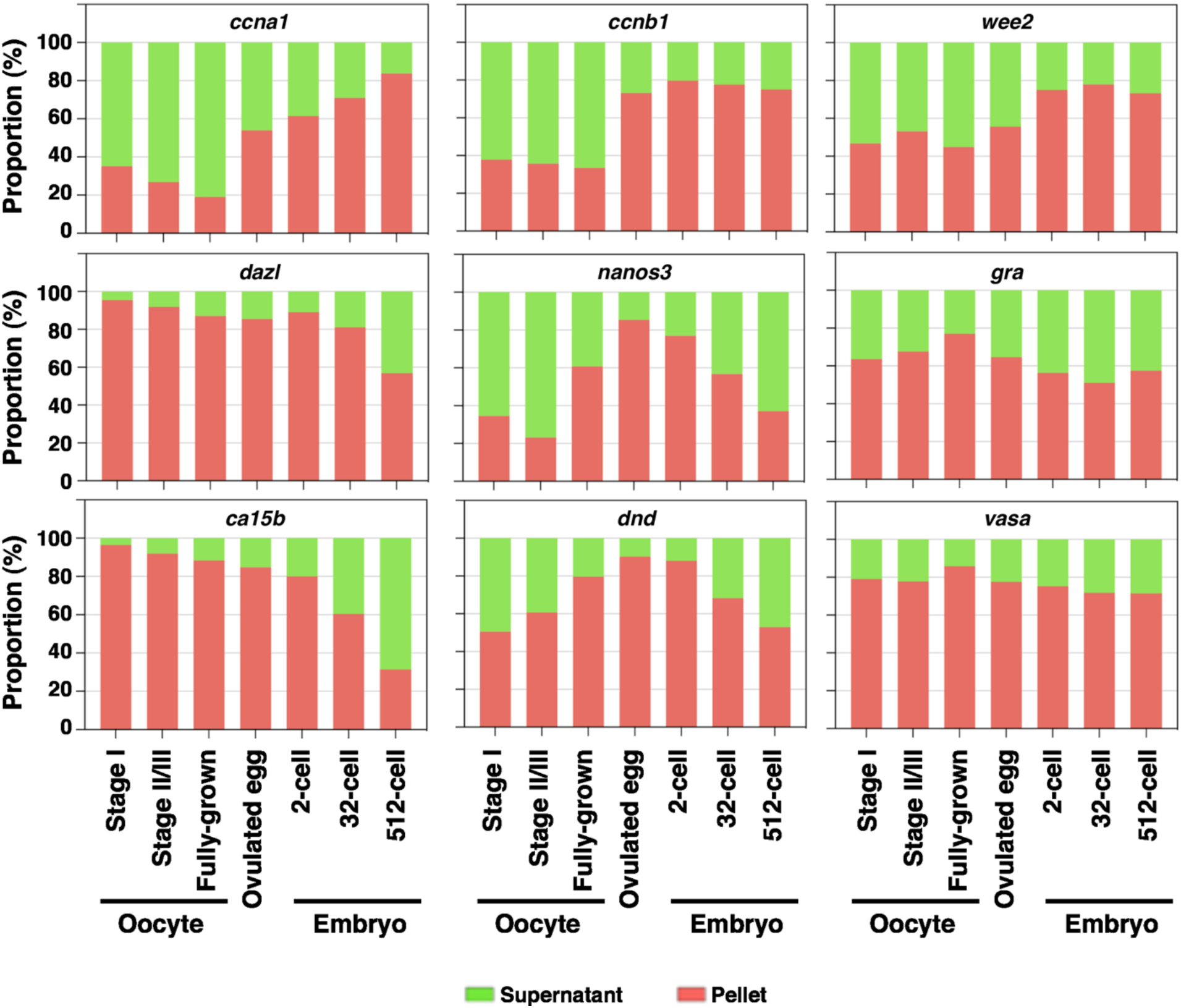
RNA phase transition during zebrafish OET. Fractionation RT-qPCR was performed to measure the percentage distribution of *ccna1, ccnb1, wee2, dazl, nanos3, gra, ca15b, dnd*, and *vasa* in stage I, stage II/III, and fully-grown oocytes, ovulated eggs, and embryos at 2-, 32-, and 512-cell stages.

## Discussion

The OET is one of the most dramatic developmental transitions during which the oocyte undergoes meiotic maturation, fuses with the sperm to form the zygote, and ultimately activates the zygotic genome to direct embryonic development. These events occur in the absence of transcription and are regulated precisely by maternal products that are synthesized and stored in the oocyte during oogenesis. Currently, it remains largely unclear how maternal products are remodeled during the OET to orchestrate the beginning of embryogenesis.

Here we report the phase transition of maternal RNA during the OET, a novel remodeling event that appears to be evolutionarily conserved. We found that during *Xenopus* oocyte maturation, a group of maternal RNAs become increasingly resistant to detergent extraction. The majority of these RNAs are asymmetrically localized in the animal hemisphere. Many of them encode cell cycle regulators that are rapidly degraded during the MZT. We tested some of these maternal genes in zebrafish. Indeed, we found that *ccna1* and *ccnb1*, which encode cyclin proteins, undergo a soluble-to-insoluble phase transition. Like in the *Xenopus* oocyte, zebrafish *ccnb1* is localized in the animal pole ^45^, suggesting that the soluble-to-insoluble phase transition of maternal mRNA is regulated by an evolutionarily conserved mechanism. In contrast to these S-I RNAs, many vegetally localized RNAs are highly insoluble in *Xenopus* oocytes, but become solubilized after oocyte maturation. Intriguingly, a subset of I-S RNAs encode proteins essential for *Xenopus* germline development. Our results reveal that some zebrafish germline RNAs are I-S transcripts as well. Interestingly, we noticed two differences between *Xenopus* and zebrafish. First of all, the phase transition of zebrafish germline RNAs occurs after fertilization, not during oocyte maturation. Secondly, some zebrafish germline RNAs, including *vasa* and *gra*, remain insoluble even at the MBT. Thus, while the phase transition of maternal RNA happens in both *Xenopus* and zebrafish, some species-specific differences clearly exist.

Our results reveal that the transition of *Xenopus* germline RNAs into the insoluble phase during early oogenesis is caused by the recruitment of germline RNAs into the germ plasm. Likewise, the solubilization of germline RNAs is a consequence of the remodeling of the germ plasm during the OET. In support of this idea, we found that Xvelo1/Buc, the matrix protein of the germ plasm ^21–24^, is essential for maintaining germline RNAs in an insoluble state. In zebrafish *bucky ball* mutant oocytes, which cannot assemble Bb/germ plasm, germline RNAs become much more soluble. During *Xenopus* oocyte maturation, Xvelo1 is markedly down-regulated, leading to the solubilization of germline RNAs. We found the solubilization of germline RNAs can be inhibited by overexpression of Xvelo1 during *Xenopus* oocyte maturation. Interestingly, the solubilization of zebrafish germline RNAs occurs after fertilization. This observation is consistent with the finding that down-regulation of Buc occurs during zebrafish early embryonic development. In a study by Riemer et al., abundant Buc was detected in the entire blastodisc after fertilization. Buc declines rapidly during cleavage. By the 128-cell and dome stage, only four Buc protein aggregates are maintained in the embryo ^49^. This explains why the timing of the solubilization of germline RNAs is different in *Xenopus* and zebrafish.

To understand the biological significance of RNA phase transition during the OET, we examined the expression level of proteins encoded by S-I, I-S, and I-I RNAs during *Xenopus* OET, using the proteomic data published recently by Peuchen et al ^50^. We failed to find any obvious correlation between RNA phase transition and changes in the expression of corresponding proteins (S-Fig 3). It seems unlikely that phase transition may serve as an important mechanism to regulate the translation of maternal RNAs at a global level during the OET.

In the case of germline RNAs, a large body of studies has demonstrated that germ plasm functions as the cargo for the vegetal transportation of germline RNAs. Hence insolubilization of germline RNAs during early oogenesis, i.e., recruitment of germline RNAs into the Bb or germ plasm, is essential for the asymmetric localization of these RNAs ^51–55^. What is the function of the phase transition of germline RNAs during the OET? Our results presented here demonstrate that the phase transition of germline RNAs influences their susceptibility to RNA degradation machinery. We show that insoluble germline RNAs are more resistant to RNase A treatment *in vitro*. In *bucky ball* mutants, where the solubility of germline RNAs is increased, a subset of germline RNAs is down-regulated. Thus, maintaining germline RNAs in an insoluble form ensures that germline RNAs are stored in a protective environment. It is well-known that after fertilization, only a fraction of germline RNAs coalesce into large aggregates and are inherited by PGCs. The remaining ones are degraded in somatic cells. Interfering with somatic clearance of germline RNAs impairs the segregation of the germline from the soma. It has been reported that overexpression of Buc prevents the degradation of germline RNAs in somatic tissue, converting somatic cells into PGCs ^22, 41^. Based on the literature and results presented here, we thus propose that the solubilization of germline RNAs during the OET, by increasing their accessibility to RNA degradation machinery, facilitates the clearance of germline RNAs in the soma.

Our finding that a fairly large number of maternal RNAs are insolubilized during oogenesis (I-S and I-I transcripts) and the OET (S-I transcripts) is fundamentally important. It argues for the existence of novel remodeling mechanisms that act on RNAs to regulate oogenesis and the OET. Apart from germline RNAs, which represent only a small portion of I-S transcripts, it is unclear how somatic I-S RNAs are insolubilized. Further work is needed to understand the biological significance of these phase transition events. Strikingly, we found the majority of S-I transcripts are rapidly degraded after zygotic genome activation. It will be of great interest to determine if there is a causal relationship between the phase transition of S-I transcripts during the OET and their degradation during the MZT.

In summary, this work demonstrates for the first time that many RNAs can transition between the soluble and insoluble phases under physiological conditions. It raises the striking possibility that as a novel post-transcriptional regulatory mechanism, precisely regulated RNA phase transition may play fundamentally important roles in a wide variety of biological processes.

## Acknowledgment

We thank Dr. Mary Mullins for providing the *bucky ball* mutant zebrafish, Jia Fu for technical support. JY is supported by a grant from NIH (R35 GM131810). WM is supported by NIH grants (R03AI146900 and R01GM140306).

## Author Contributions

Study conception and design: JY.

Data collection: HH, SC, MM, D, HF, EB, and JY.

Analysis and interpretation of results: HH, SC, WM, EB, and JY.

Draft manuscript preparation: HH and JY.

All authors reviewed the results and approved the final version of the manuscript.

## STAR Methods

### Resource Availability

#### Lead contact

- Further information and requests for resources and reagents should be directed to and will be fulfilled by the lead contact, Jing Yang.

#### Materials availability

- This study did not generate any new unique reagents.

#### Data and code availability

- RNA sequencing data analyzed in this study have been deposited to the Gene Expression Omnibus (GEO) with the dataset identifier GSE199254.
- This paper does not report the original code.
- Any additional information required to reanalyze the data reported in this paper is available from the lead contact upon request.

### Experimental Model and Subject Details

#### Xenopus laevis

*Xenopus* procedures were approved by the University of Illinois at Urbana-Champaign Institutional Animal Care and Use Committee (IACUC) under animal protocol #20125 and performed in accordance with the recommendations of the Guide for the Care and Use of Laboratory Animals of the National Institutes of Health. Oocytes were collected from ovarian tissues by manual defolliculation or collagenase treatment and cultured in the oocyte culture medium (OCM) ^56^. In order to induce oocyte meiotic maturation, stage VI oocytes were cultured in the OCM containing 2 µM progesterone and incubated at 18 °C overnight. Different stages of oocytes and mature eggs were collected for further analysis. For microinjection, 2 ng *xvelo1* mRNAs were injected into the oocyte using a Narishige IM300 microinjector, and then the oocyte was either cultured in the OCM or treated with progesterone.

#### Danio rerio

Zebrafish procedures were approved by the University of Illinois at Urbana-Champaign Institutional Animal Care and Use Committee (IACUC) under animal protocol #21179 and performed in accordance with the recommendations of the Guide for the Care and Use of Laboratory Animals of the National Institutes of Health. The female Tübingen strain was purchased from ZIRC (Zebrafish International Resource Center), and Buckyball mutant zebrafish were received from Dr. Mullins’ lab at the University of Pennsylvania. Zebrafish oocytes were obtained as described ^57^. Briefly, ovarian tissues were dissected and dissociated by 15 mg/ml collagenase treatment for 30 min at room temperature (RT). Then, each stage of oocytes was sorted and collected manually for further experiments.

### Method Details

#### Cell Fractionation

Fractionation was performed as described ^16, 58^ with slight modification. Briefly, *Xenopus* oocytes and mature eggs were rinsed with ice-cold washing buffer (150 mM KOAc, 2.5 mM MgOAc_2_, and 20 mM K-HEPES pH 7.4). To extract soluble fraction, four oocytes or eggs were homogenized in 160 µl of ice-cold extraction buffer (150 mM KOAc, 2.5 mM MgOAc_2_, 20 mM K-HEPES pH 7.4, 2 mM DTT, 1 mM PSMF, 50 µg/ml cycloheximide, and 200 units/ml RNase inhibitor) containing 0.5 % Triton X-100 and centrifuged at 800 x g for 5 min. After centrifuging, supernatants were transferred to new tubes and centrifuged again at 10,000 x g for 10 min to remove contaminating organelles and cell debris. Pellets were resuspended in 160 µl of ice-cold extraction buffer with 0.5 % Triton X-100. Twice centrifuged supernatants as soluble fractions and resuspended pellet solution as insoluble fractions were prepared for further analysis. For the fractionation of zebrafish samples, five samples of each zebrafish oocytes, ovulated eggs, and embryos were homogenized in 100 µl of ice-cold extraction buffer containing 0.5 % Triton X-100 and centrifuged at 10,000 x g for 10 min. After centrifuging, supernatants were transferred to new tubes, and pellets were resuspended with 100 µl of ice-cold extraction buffer containing 0.5 % Triton X-100. All centrifugations were performed in a refrigerated centrifuge at 4 °C.

#### Whole-mount immunofluorescence

*Xenopus* stage VI and mature eggs were fixed in Dent’s fixative (80 % of methanol and 20 % DMSO) at −20 °C overnight. After fixation, samples were washed with 100 % methanol three times for 5 min each time and stored in 100 % methanol at −20 °C until the staining procedure was initiated. For whole-mount immunofluorescence, samples were rehydrated in serial dilution of methanol with TBST (TBS with 0.1% Triton X-100) for 5 min each at RT. Oocytes and eggs were bisected by a razor blade to obtain hemi-sectioned and vegetal hemisphere samples. After bisection, samples were washed in TBST once and incubated in blocking buffer (0.15 % Triton X-100, 2 % BSA in TBS) containing 10 % normal serum from the same host as the secondary antibody for an hour at RT. Then, samples were incubated in blocking buffer with the primary antibodies (1:100) overnight at 4 °C. Samples were washed with TBST six times for 30 min each, incubated in blocking buffer with the secondary antibodies (1:500) overnight at 4 °C, and washed with TBST six times for 30 min each. Stained samples were washed with 100 % methanol twice for 10 min each and then added in BABB (1 : 2 ratio of benzyl alcohol : benzyl benzoate) to be cleared. Images were acquired by Nikon A1Rsi confocal microscope.

#### RNase A assay

For time-dependent RNase A assay, 40 oocytes and 40 mature eggs were harvested and homogenized in 800 µl of lysis buffer. 100 µl of the lysates was saved as ‘no RNase A’. The remaining lysates were transferred into six tubes, each containing 100 µl, and treated with RNase A (12.5 pg/µl; final concentration) at 37 °C for 5, 15, 30, 60, 90, and 120 min. After the RNase A treatment, RNAs were extracted for RT-qPCR. For fractionation RNase A assay, 20 oocytes and 20 mature eggs were harvested and homogenized in 800 µl of extraction buffer (150 mM KOAc, 2.5 mM MgOAc_2_, 20 mM K-HEPES pH 7.4, 2 mM DTT, 1 mM PSMF, 50 µg/ml cycloheximide, and 200 units/ml RNase inhibitor) containing 0.5 % Triton X-100. 160 µl of the lysates was saved as ‘no RNase A’. The remaining lysates were transferred into 4 tubes (160 µl each) and incubated in different concentrations (1 pg/µl, 10 pg/µl, 100 pg/µl, and 1 ng/µl) of RNase A at 37 °C for 2 hr. After incubation with RNase A, samples were fractionated into soluble and insoluble fractions by the above protocol. RNAs were extracted using TRIzol reagent according to the manufacturer’s instructions.

### *In vitro* transcription

*Xvelo1* mRNAs were synthesized from 2 µg plasmid templates using the mMESSAGE mMACHINE Kit for SP6. All probes used in this study were synthesized from 2 µg plasmid templates using T3 RNA polymerase. To be incorporated Digoxigenin (DIG) UTPs into the probes, 1 µl of DIG RNA labeling mix was added to the *in vitro* transcription reaction solution. Dig-labeled probes were detected by anti-Digoxigenin-AP Fab fragments/BM-purple staining.

### Xenopus in situ hybridization

Oocytes at various stages were fixed with MEMFA (0.1 M MOPS pH 7.4, 2 mM EGTA, 1 mM MgSO_4_, and 3.7% formaldehyde solution) for an hour at RT, washed with PBS twice, and dehydrated in methanol. Dehydrated samples were stored in 100 % methanol at −20°C. For *in situ* hybridization, all samples were rehydrated in serial dilution of methanol with PBSW (PBS with 0.1 % Tween-20) for 5 min each at RT. Stage III, IV, V, and VI oocytes were hemi-sectioned by a razor blade. Once all stage II oocytes and bisected mid and late-stage oocytes were prepared, *in situ* hybridization was performed as described ^59^.

### RNA Extraction and Quantitative RT-PCR

RNAs were extracted from the soluble and insoluble fraction of oocytes, eggs, or embryos using TRIzol reagent in accordance with the manufacturer’s instructions. Then, reverse transcription and real-time PCR were performed according to standard protocols to analyze the expression level of mRNAs. Ct values were acquired by Applied Biosystems QuantStudio 3 Real-Time PCR System. All primers used are listed in the supplementary table.

### Immunoprecipitation and Western Blots

For immunoprecipitation, fully-grown oocytes were homogenized in 0.5 % NP-40 lysis buffer (50 mM Tris pH 7.6, 125mM NaCl, 1 mM EDTA, 0.5 % NP-40) with protease inhibitor cocktails and centrifuged at 20,000 x g for 10 min. Then, supernatants were transferred to new tubes. Xvelo1 antibodies were added to the supernatant and incubated overnight at 4 °C. Next, protein G-coupled agarose beads were added to precipitate Xvelo1 proteins. After washing the beads with lysis buffer three times (5 min each), proteins were eluted in 2 x SDS sample buffer by boiling for 5 min at 100°C. For western blots, fractionated samples as lysates were mixed with 2 x SDS sample buffer and boiled for 5 min at 100°C. The lysates for IP or western blots were loaded on SDS-PAGE and transferred to PVDF membranes for western blotting according to the standard protocol for western blots.

### Mitochondria and endoplasmic reticulum (ER) staining of Zebrafish oocyte

Mitochondria and ER staining were performed as described ^57^ with slight modification. Zebrafish stage I oocytes were isolated from the ovarian tissues of the wild-type and Buc mutant zebrafish according to the protocol mentioned above. Stage I oocytes were stained with 0.5 µM of MitoTracker and 0.5 µM of ER tracker each in staining buffer (PBS with 0.1 % BSA) at RT for 30 min. Then, samples were washed with staining buffer three times for 10 min each at RT. After washing, samples were stained with 0.5 µg/ml of Hoechst 33342 in the staining buffer for 10 min at RT and then mounted with staining buffer in glass concavity slides. All images were acquired using Nikon A1Rsi confocal microscope.

### Analysis of RNA-Sequencing

All RNA-seq datasets used in this study were listed in the Key resources table. The fractionation RNA-seq was analyzed as described ^16^ with slight modification. Briefly, all analyses were performed using abundant transcripts (17,811) and a proportion scale according to the previous research analysis ^16^. The soluble fraction is the sum of the cytosolic and ER fractions, and the pellet was considered insoluble. These 17,811 transcripts were classified into three groups below.

1. The Soluble-to-Insoluble Phase Transition (S-I): Transcripts that were increased more than two times and were enriched more than 15% in the pellet fraction of mature eggs after oocyte maturation (863).
2. The Insoluble-to-Soluble Phase Transition (I-S): Transcripts that were enriched more than 40% in the pellet fraction of oocytes and those pellet fraction is reduced by more than 25% after oocyte maturation (165).
3. The Insoluble-Insoluble State (I-I): Among the transcripts enriched more than 40% in the oocyte pellet fraction, the rest of the transcripts excluding those belonging to the I-S (68).

All GO and Protein-Protein interaction (PPI) results were generated by the criteria above.

#### Comparison to RNA localization data along the A-V axis in *Xenopus* eggs

The distribution of 15,005 transcripts along the animal-vegetal axis in the *Xenopus laevis* eggs has been reported previously ^26^. 10,798 transcripts were detected in both our fractionation RNA-seq and the Sindelka study. Among 10,798 transcripts, 573, 88, and 4 transcripts were detected in S-I, I-S, and I-I groups, respectively. The heat maps were generated by the expression level of each section along the animal-vegetal axis.

#### Comparison to RNA expression level during the early *Xenopus* embryonic development

RNA-seq dataset analyzing the expression level of *Xenopus* early embryos was reported previously ^31^. 11,169 transcripts were detected in both our fractionation RNA-seq and the Session study. Among 11,169 transcripts, genes that are expressed 0 at the egg stage were filtered out (105). In this analysis, datasets for mature eggs, stages 8, 9, 10, 12, and 15 embryos were used, and the results were normalized by the expression level of mature eggs. Since this analysis aimed to show the maternal RNA degradation in early embryonic development, zygotic genome activation was not considered in the expression analysis. Therefore, when the expression level of a specific transcript was more than 1 after normalization, that gene was considered zygotic gene expression started. In this study, when the expression level of a specific gene was decreased by more than 25 % in the following stage, that gene was deemed to occurring degradation. Based on those criteria, all transcripts were classified into three groups below.

1. Class A (Rapid degraded genes, 2288 transcripts): Transcripts that started to degrade between the mature egg to stage 8 embryos and between stages 8 to stage 9 embryos.
2. Class B (Slowly degraded genes, 6101 transcripts): Transcripts that started to degrade between stages 9 to 10 embryos and between stages 10 to 11 embryos.
3. Class C (Relatively stable genes, 2675 transcripts): Transcripts that excluded those belonging to classes A and B.

All heat map results and diagrams were generated by the criteria above.

#### Comparison to proteomic data in *Xenopus* oocytes and embryos

The expression levels of proteins from stage VI oocyte to stage 2 embryo have been reported previously ^50^. Among all proteins detected by mass spec, 166, 21, and 2 were encoded by S-I, I-S, and I-I transcripts, respectively. The expression of these proteins was converted to the log scale and normalized by the expression levels in mature eggs. The heatmap was generated based on the protein expression levels in each stage to visualize the changes in protein levels.

### Quantification and Statistical Analysis

All information about the statistical details is provided in the figure legends. Visualization for RNA-seq and proteomic profile-related results was performed by R studio. All image analysis and statistical tests were performed by ImageJ ^60^ and GraphPad Prism 9, respectively.

## Key resources table

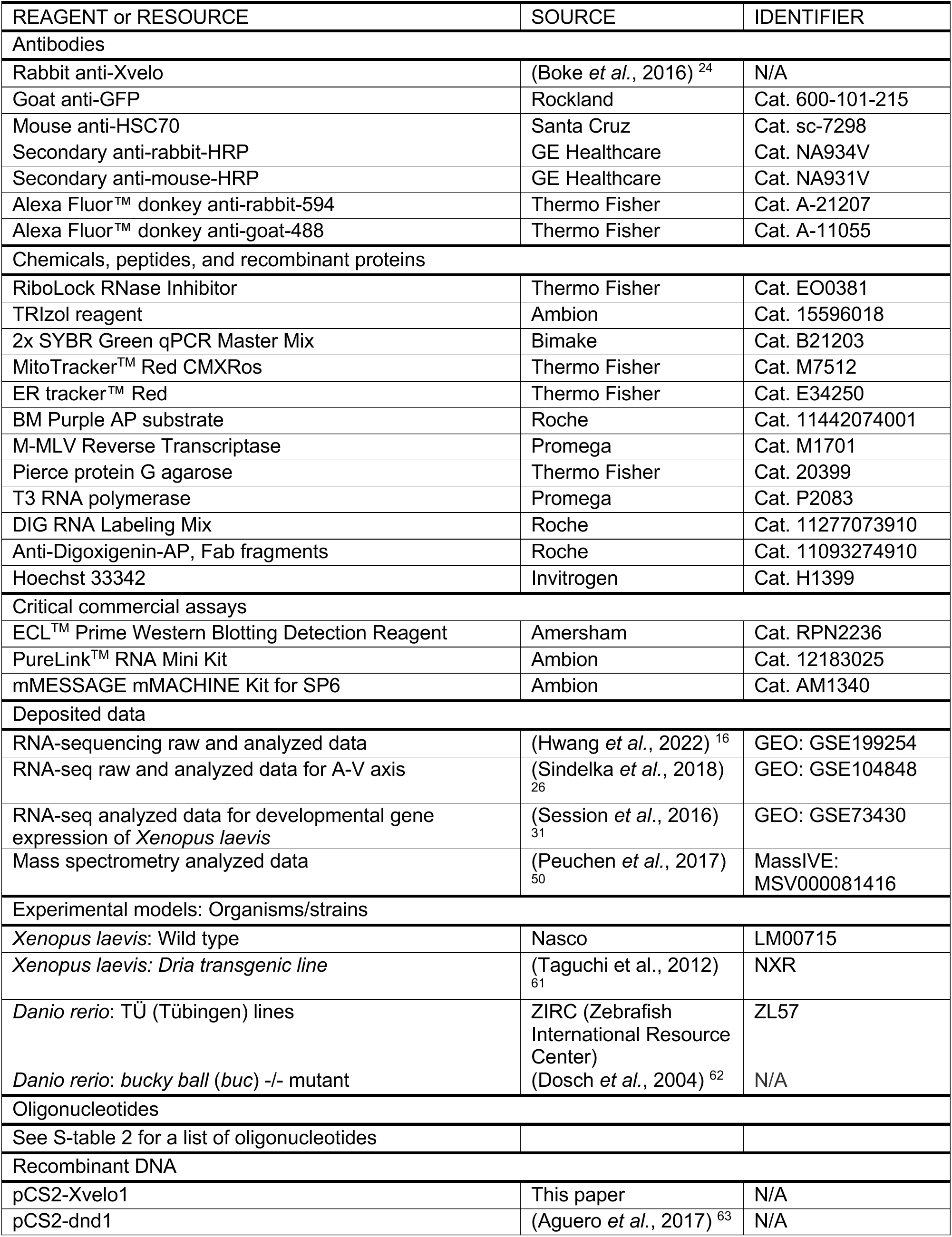
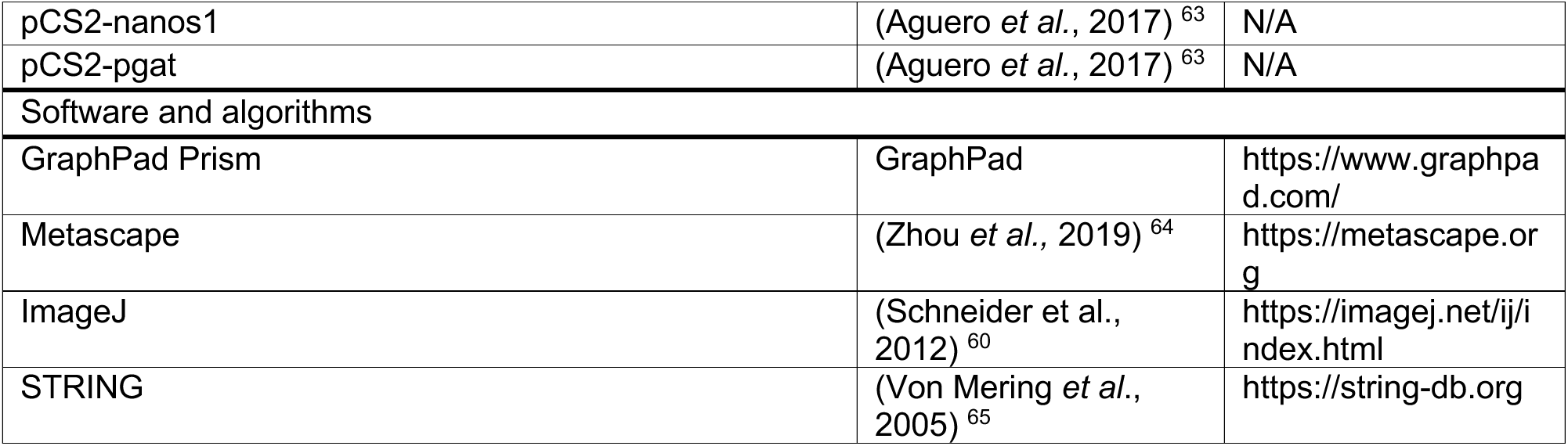

| REAGENT or RESOURCE | SOURCE | IDENTIFIER |
| --- | --- | --- |
| Antibodies |  |  |
| Rabbit anti-Xvelo | (Boke *et al.*, 2016)<sup>24</sup> | N/A |
| Goat anti-GFP | Rockland | Cat. 600-101-215 |
| Mouse anti-HSC70 | Santa Cruz | Cat. sc-7298 |
| Secondary anti-rabbit-HRP | GE Healthcare | Cat. NA934V |
| Secondary anti-mouse-HRP | GE Healthcare | Cat. NA931V |
| Alexa Fluor™ donkey anti-rabbit-594 | Thermo Fisher | Cat. A-21207 |
| Alexa Fluor™ donkey anti-goat-488 | Thermo Fisher | Cat. A-11055 |
| Chemicals, peptides, and recombinant proteins |  |  |
| RiboLock RNase Inhibitor | Thermo Fisher | Cat. EO0381 |
| TRIzol reagent | Ambion | Cat. 15596018 |
| 2x SYBR Green qPCR Master Mix | Bimake | Cat. B21203 |
| MitoTracker™ Red CMXRos | Thermo Fisher | Cat. M7512 |
| ER tracker™ Red | Thermo Fisher | Cat. E34250 |
| BM Purple AP substrate | Roche | Cat. 11442074001 |
| M-MLV Reverse Transcriptase | Promega | Cat. M1701 |
| Pierce protein G agarose | Thermo Fisher | Cat. 20399 |
| T3 RNA polymerase | Promega | Cat. P2083 |
| DIG RNA Labeling Mix | Roche | Cat. 11277073910 |
| Anti-Digoxigenin-AP, Fab fragments | Roche | Cat. 11093274910 |
| Hoechst 33342 | Invitrogen | Cat. H1399 |
| Critical commercial assays |  |  |
| ECL™ Prime Western Blotting Detection Reagent | Amersham | Cat. RPN2236 |
| PureLink™ RNA Mini Kit | Ambion | Cat. 12183025 |
| mMESSAGE mMACHINE Kit for SP6 | Ambion | Cat. AM1340 |
| Deposited data |  |  |
| RNA-sequencing raw and analyzed data | (Hwang *et al.*, 2022)<sup>16</sup> | GEO: GSE199254 |
| RNA-seq raw and analyzed data for A-V axis | (Sindelka *et al.*, 2018)<sup>26</sup> | GEO: GSE104848 |
| RNA-seq analyzed data for developmental gene expression of *Xenopus laevis* | (Session *et al.*, 2016)<sup>31</sup> | GEO: GSE73430 |
| Mass spectrometry analyzed data | (Peuchen *et al.*, 2017)<sup>50</sup> | MassIVE: MSV000081416 |
| Experimental models: Organisms/strains |  |  |
| *Xenopus laevis*: Wild type | Nasco | LM00715 |
| *Xenopus laevis*: *Dria transgenic line* | (Taguchi *et al.*, 2012)<sup>61</sup> | NXR |
| *Danio rerio*: Tü (Tübingen) lines | ZIRC (Zebrafish International Resource Center) | ZL57 |
| *Danio rerio*: *bucky ball (buc)* -/- mutant | (Dosch *et al.*, 2004)<sup>62</sup> | N/A |
| Oligonucleotides |  |  |
| See S-table 2 for a list of oligonucleotides |  |  |
| Recombinant DNA |  |  |
| pCS2-Xvelo1 | This paper | N/A |
| pCS2-dnd1 | (Aguero *et al.*, 2017)<sup>63</sup> | N/A |
| pCS2-nanos1 | (Aguero *et al.*, 2017)<sup>63</sup> | N/A |
| pCS2-pgat | (Aguero *et al.*, 2017)<sup>63</sup> | N/A |
| Software and algorithms |  |  |
| GraphPad Prism | GraphPad | https://www.graphpad.com/ |
| Metascape | (Zhou *et al.*, 2019)<sup>64</sup> | https://metascape.org |
| ImageJ | (Schneider *et al.*, 2012)<sup>60</sup> | https://imagej.net/ij/index.html |
| STRING | (Von Mering *et al.*, 2005)<sup>65</sup> | https://string-db.org |

## References

1. Schultz, R.M., Stein, P., and Svoboda, P. (2018). The oocyte-to-embryo transition in mouse: past, present, and future. Biol Reprod 99, 160–174. 10.1093/biolre/ioy013.

2. Jaffe, L.A., and Terasaki, M. (1994). Structural changes in the endoplasmic reticulum of starfish oocytes during meiotic maturation and fertilization. Dev Biol 164, 579–587. 10.1006/dbio.1994.1225.

3. Terasaki, M., and Jaffe, L.A. (1991). Organization of the sea urchin egg endoplasmic reticulum and its reorganization at fertilization. J Cell Biol 114, 929–940.

4. Mehlmann, L.M., Terasaki, M., Jaffe, L.A., and Kline, D. (1995). Reorganization of the endoplasmic reticulum during meiotic maturation of the mouse oocyte. Dev Biol 170, 607–615. 10.1006/dbio.1995.1240.

5. Shiraishi, K., Okada, A., Shirakawa, H., Nakanishi, S., Mikoshiba, K., and Miyazaki, S. (1995). Developmental changes in the distribution of the endoplasmic reticulum and inositol 1,4,5-trisphosphate receptors and the spatial pattern of Ca2+ release during maturation of hamster oocytes. Dev Biol 170, 594–606. 10.1006/dbio.1995.1239.

6. Kume, S., Yamamoto, A., Inoue, T., Muto, A., Okano, H., and Mikoshiba, K. (1997). Developmental expression of the inositol 1,4,5-trisphosphate receptor and structural changes in the endoplasmic reticulum during oogenesis and meiotic maturation of Xenopus laevis. Dev Biol 182, 228–239. 10.1006/dbio.1996.8479.

7. Kline, D. (2000). Attributes and dynamics of the endoplasmic reticulum in mammalian eggs. Current topics in developmental biology 50, 125–154.

8. Terasaki, M., Runft, L.L., and Hand, A.R. (2001). Changes in organization of the endoplasmic reticulum during Xenopus oocyte maturation and activation. Mol Biol Cell 12, 1103–1116. 10.1091/mbc.12.4.1103.

9. FitzHarris, G., Marangos, P., and Carroll, J. (2007). Changes in endoplasmic reticulum structure during mouse oocyte maturation are controlled by the cytoskeleton and cytoplasmic dynein. Dev Biol 305, 133–144. 10.1016/j.ydbio.2007.02.006.

10. Stitzel, M.L., and Seydoux, G. (2007). Regulation of the oocyte-to-zygote transition. Science 316, 407–408. 10.1126/science.1138236.

11. Deshler, J.O., Highett, M.I., and Schnapp, B.J. (1997). Localization of Xenopus Vg1 mRNA by Vera protein and the endoplasmic reticulum. Science 276, 1128–1131. 10.1126/science.276.5315.1128.

12. Alarcon, V.B., and Elinson, R.P. (2001). RNA anchoring in the vegetal cortex of the Xenopus oocyte. J Cell Sci 114, 1731–1741.

13. Chang, P., Torres, J., Lewis, R.A., Mowry, K.L., Houliston, E., and King, M.L. (2004). Localization of RNAs to the mitochondrial cloud in Xenopus oocytes through entrapment and association with endoplasmic reticulum. Mol Biol Cell 15, 4669–4681. 10.1091/mbc.e04-03-0265.

14. Prodon, F., Dru, P., Roegiers, F., and Sardet, C. (2005). Polarity of the ascidian egg cortex and relocalization of cER and mRNAs in the early embryo. J Cell Sci 118, 2393–2404. 10.1242/jcs.02366.

15. Sardet, C., Nishida, H., Prodon, F., and Sawada, K. (2003). Maternal mRNAs of PEM and macho 1, the ascidian muscle determinant, associate and move with a rough endoplasmic reticulum network in the egg cortex. Development 130, 5839–5849. 10.1242/dev.00805.

16. Hwang, H., Yun, S., Arcanjo, R.B., Divyanshi, Chen, S., Mei, W., Nowak, R.A., Kwon, T., and Yang, J. (2022). Regulation of RNA localization during oocyte maturation by dynamic RNA-ER association and remodeling of the ER. Cell Rep 41, 111802. 10.1016/j.celrep.2022.111802.

17. Solter, D., Hiiragi, T., Evsikov, A.V., Moyer, J., De Vries, W.N., Peaston, A.E., and Knowles, B.B. (2004). Epigenetic mechanisms in early mammalian development. Cold Spring Harb Symp Quant Biol 69, 11–17. 10.1101/sqb.2004.69.11.

18. Hwang, H., Jin, Z., Krishnamurthy, V.V., Saha, A., Klein, P.S., Garcia, B., Mei, W., King, M.L., Zhang, K., and Yang, J. (2019). Novel functions of the ubiquitin-independent proteasome system in regulating Xenopus germline development. Development 146. 10.1242/dev.172700.

19. Liu, Y., Zhao, H., Shao, F., Zhang, Y., Nie, H., Zhang, J., Li, C., Hou, Z., Chen, Z.J., Wang, J., et al. (2023). Remodeling of maternal mRNA through poly(A) tail orchestrates human oocyte-to-embryo transition. Nat Struct Mol Biol. 10.1038/s41594-022-00908-2.

20. Shi, B., Zhang, J., Heng, J., Gong, J., Zhang, T., Li, P., Sun, B.F., Yang, Y., Zhang, N., Zhao, Y.L., et al. (2020). RNA structural dynamics regulate early embryogenesis through controlling transcriptome fate and function. Genome Biol 21, 120. 10.1186/s13059-020-02022-2.

21. Marlow, F.L., and Mullins, M.C. (2008). Bucky ball functions in Balbiani body assembly and animal-vegetal polarity in the oocyte and follicle cell layer in zebrafish. Dev Biol 321, 40–50. 10.1016/j.ydbio.2008.05.557.

22. Bontems, F., Stein, A., Marlow, F., Lyautey, J., Gupta, T., Mullins, M.C., and Dosch, R. (2009). Bucky ball organizes germ plasm assembly in zebrafish. Curr Biol 19, 414–422. 10.1016/j.cub.2009.01.038.

23. Nijjar, S., and Woodland, H.R. (2013). Protein interactions in Xenopus germ plasm RNP particles. PLoS ONE 8, e80077. 10.1371/journal.pone.0080077.

24. Boke, E., Ruer, M., Wuhr, M., Coughlin, M., Lemaitre, R., Gygi, S.P., Alberti, S., Drechsel, D., Hyman, A.A., and Mitchison, T.J. (2016). Amyloid-like Self-Assembly of a Cellular Compartment. Cell 166, 637–650. 10.1016/j.cell.2016.06.051.

25. Houston, D.W. (2013). Regulation of cell polarity and RNA localization in vertebrate oocytes. Int Rev Cell Mol Biol 306, 127–185. 10.1016/B978-0-12-407694-5.00004-3.

26. Sindelka, R., Abaffy, P., Qu, Y., Tomankova, S., Sidova, M., Naraine, R., Kolar, M., Peuchen, E., Sun, L., Dovichi, N., and Kubista, M. (2018). Asymmetric distribution of biomolecules of maternal origin in the Xenopus laevis egg and their impact on the developmental plan. Sci Rep 8, 8315. 10.1038/s41598-018-26592-1.

27. Newport, J., and Kirschner, M. (1982). A major developmental transition in early Xenopus embryos: II. Control of the onset of transcription. Cell 30, 687–696. 0092-8674(82)90273-2 [pii].

28. Newport, J., and Kirschner, M. (1982). A major developmental transition in early Xenopus embryos: I. characterization and timing of cellular changes at the midblastula stage. Cell 30, 675–686. 0092-8674(82)90272-0 [pii].

29. Howe, J.A., Howell, M., Hunt, T., and Newport, J.W. (1995). Identification of a developmental timer regulating the stability of embryonic cyclin A and a new somatic A-type cyclin at gastrulation. Genes Dev 9, 1164–1176.

30. Howe, J.A., and Newport, J.W. (1996). A developmental timer regulates degradation of cyclin E1 at the midblastula transition during Xenopus embryogenesis. Proc Natl Acad Sci U S A 93, 2060–2064. 10.1073/pnas.93.5.2060.

31. Session, A.M., Uno, Y., Kwon, T., Chapman, J.A., Toyoda, A., Takahashi, S., Fukui, A., Hikosaka, A., Suzuki, A., Kondo, M., et al. (2016). Genome evolution in the allotetraploid frog Xenopus laevis. Nature 538, 336–343. 10.1038/nature19840.

32. MacArthur, H., Houston, D.W., Bubunenko, M., Mosquera, L., and King, M.L. (2000). DEADSouth is a germ plasm specific DEAD-box RNA helicase in Xenopus related to eIF4A. Mechanisms of development 95, 291–295.

33. Zhou, Y., and King, M.L. (1996). Localization of Xcat-2 RNA, a putative germ plasm component, to the mitochondrial cloud in Xenopus stage I oocytes. Development 122, 2947–2953.

34. Horvay, K., Claussen, M., Katzer, M., Landgrebe, J., and Pieler, T. (2006). Xenopus Dead end mRNA is a localized maternal determinant that serves a conserved function in germ cell development. Dev Biol 291, 1–11. 10.1016/j.ydbio.2005.06.013.

35. Owens, D.A., Butler, A.M., Aguero, T.H., Newman, K.M., Van Booven, D., and King, M.L. (2017). High-throughput analysis reveals novel maternal germline RNAs crucial for primordial germ cell preservation and proper migration. Development 144, 292–304. 10.1242/dev.139220.

36. Houston, D.W., Zhang, J., Maines, J.Z., Wasserman, S.A., and King, M.L. (1998). A Xenopus DAZ-like gene encodes an RNA component of germ plasm and is a functional homologue of Drosophila boule. Development 125, 171–180.

37. Hudson, C., and Woodland, H.R. (1998). Xpat, a gene expressed specifically in germ plasm and primordial germ cells of Xenopus laevis. Mechanisms of development 73, 159–168.

38. Tarbashevich, K., Koebernick, K., and Pieler, T. (2007). XGRIP2.1 is encoded by a vegetally localizing, maternal mRNA and functions in germ cell development and anteroposterior PGC positioning in Xenopus laevis. Dev Biol 311, 554–565. 10.1016/j.ydbio.2007.09.012.

39. Oh, D., and Houston, D.W. (2017). Role of maternal Xenopus syntabulin in germ plasm aggregation and primordial germ cell specification. Dev Biol 432, 237–247. 10.1016/j.ydbio.2017.10.006.

40. Claussen, M., and Pieler, T. (2004). Xvelo1 uses a novel 75-nucleotide signal sequence that drives vegetal localization along the late pathway in Xenopus oocytes. Dev Biol 266, 270–284. 10.1016/j.ydbio.2003.09.043.

41. Ye, D., Zhu, L., Zhang, Q., Xiong, F., Wang, H., Wang, X., He, M., Zhu, Z., and Sun, Y. (2019). Abundance of Early Embryonic Primordial Germ Cells Promotes Zebrafish Female Differentiation as Revealed by Lifetime Labeling of Germline. Mar Biotechnol (NY) 21, 217–228. 10.1007/s10126-019-09874-1.

42. Elkouby, Y.M., Jamieson-Lucy, A., and Mullins, M.C. (2016). Oocyte Polarization Is Coupled to the Chromosomal Bouquet, a Conserved Polarized Nuclear Configuration in Meiosis. PLoS Biol 14, e1002335. 10.1371/journal.pbio.1002335.

43. Beer, R.L., and Draper, B.W. (2013). nanos3 maintains germline stem cells and expression of the conserved germline stem cell gene nanos2 in the zebrafish ovary. Dev Biol 374, 308–318. 10.1016/j.ydbio.2012.12.003.

44. Weidinger, G., Stebler, J., Slanchev, K., Dumstrei, K., Wise, C., Lovell-Badge, R., Thisse, C., Thisse, B., and Raz, E. (2003). dead end, a novel vertebrate germ plasm component, is required for zebrafish primordial germ cell migration and survival. Curr Biol 13, 1429–1434.

45. Howley, C., and Ho, R.K. (2000). mRNA localization patterns in zebrafish oocytes. Mechanisms of development 92, 305–309. 10.1016/s0925-4773(00)00247-1.

46. Wang, H., Teng, Y., Xie, Y., Wang, B., Leng, Y., Shu, H., and Deng, F. (2013). Characterization of the carbonic anhydrases 15b expressed in PGCs during early zebrafish development. Theriogenology 79, 443–452. 10.1016/j.theriogenology.2012.10.016.

47. Strasser, M.J., Mackenzie, N.C., Dumstrei, K., Nakkrasae, L.I., Stebler, J., and Raz, E. (2008). Control over the morphology and segregation of Zebrafish germ cell granules during embryonic development. BMC Dev Biol 8, 58. 10.1186/1471-213X-8-58.

48. Yoon, C., Kawakami, K., and Hopkins, N. (1997). Zebrafish vasa homologue RNA is localized to the cleavage planes of 2- and 4-cell-stage embryos and is expressed in the primordial germ cells. Development 124, 3157–3165. 10.1242/dev.124.16.3157.

49. Riemer, S., Bontems, F., Krishnakumar, P., Gomann, J., and Dosch, R. (2015). A functional Bucky ball-GFP transgene visualizes germ plasm in living zebrafish. Gene Expr Patterns 18, 44–52. 10.1016/j.gep.2015.05.003.

50. Peuchen, E.H., Cox, O.F., Sun, L., Hebert, A.S., Coon, J.J., Champion, M.M., Dovichi, N.J., and Huber, P.W. (2017). Phosphorylation Dynamics Dominate the Regulated Proteome during Early Xenopus Development. Sci Rep 7, 15647. 10.1038/s41598-017-15936-y.

51. Kloc, M., Bilinski, S., Chan, A.P., Allen, L.H., Zearfoss, N.R., and Etkin, L.D. (2001). RNA localization and germ cell determination in Xenopus. Int Rev Cytol 203, 63–91.

52. King, M.L., Messitt, T.J., and Mowry, K.L. (2005). Putting RNAs in the right place at the right time: RNA localization in the frog oocyte. Biol Cell 97, 19–33.

53. Oh, D., and Houston, D.W. (2017). RNA Localization in the Vertebrate Oocyte: Establishment of Oocyte Polarity and Localized mRNA Assemblages. Results Probl Cell Differ 63, 189–208. 10.1007/978-3-319-60855-6_9.

54. Jamieson-Lucy, A., and Mullins, M.C. (2019). The vertebrate Balbiani body, germ plasm, and oocyte polarity. Current topics in developmental biology 135, 1–34. 10.1016/bs.ctdb.2019.04.003.

55. O’Connell, L.C., and Mowry, K.L. (2021). Regulation of spatially restricted gene expression: linking RNA localization and phase separation. Biochem Soc Trans 49, 2591–2600. 10.1042/BST20210320.

56. Houston, D.W. (2018). Oocyte Host-Transfer and Maternal mRNA Depletion Experiments in Xenopus. Cold Spring Harb Protoc 2018. 10.1101/pdb.prot096982.

57. Lee, K.L., and Marlow, F.L. (2019). Visualizing the Balbiani Body in Zebrafish Oocytes. Methods Mol Biol 1920, 277–293. 10.1007/978-1-4939-9009-2_16.

58. Lerner, R.S., Seiser, R.M., Zheng, T., Lager, P.J., Reedy, M.C., Keene, J.D., and Nicchitta, C.V. (2003). Partitioning and translation of mRNAs encoding soluble proteins on membrane-bound ribosomes. RNA 9, 1123–1137. 10.1261/rna.5610403.

59. Rorick, A.M., Mei, W., Liette, N.L., Phiel, C., El-Hodiri, H.M., and Yang, J. (2007). PP2A:B56epsilon is required for eye induction and eye field separation. Dev Biol 302, 477–493.

60. Schneider, C.A., Rasband, W.S., and Eliceiri, K.W. (2012). NIH Image to ImageJ: 25 years of image analysis. Nat Methods 9, 671–675. 10.1038/nmeth.2089.

61. Taguchi, A., Takii, M., Motoishi, M., Orii, H., Mochii, M., and Watanabe, K. (2012). Analysis of localization and reorganization of germ plasm in Xenopus transgenic line with fluorescence-labeled mitochondria. Dev Growth Differ 54, 767–776. 10.1111/dgd.12005.

62. Dosch, R., Wagner, D.S., Mintzer, K.A., Runke, G., Wiemelt, A.P., and Mullins, M.C. (2004). Maternal control of vertebrate development before the midblastula transition: mutants from the zebrafish I. Dev Cell 6, 771–780. 10.1016/j.devcel.2004.05.002.

63. Aguero, T., Jin, Z., Chorghade, S., Kalsotra, A., King, M.L., and Yang, J. (2017). Maternal Dead-end 1 promotes translation of nanos1 by binding the eIF3 complex. Development 144, 3755–3765. 10.1242/dev.152611.

64. Zhou, Y., Zhou, B., Pache, L., Chang, M., Khodabakhshi, A.H., Tanaseichuk, O., Benner, C., and Chanda, S.K. (2019). Metascape provides a biologist-oriented resource for the analysis of systems-level datasets. Nature communications 10, 1523. 10.1038/s41467-019-09234-6.

65. von Mering, C., Jensen, L.J., Snel, B., Hooper, S.D., Krupp, M., Foglierini, M., Jouffre, N., Huynen, M.A., and Bork, P. (2005). STRING: known and predicted protein-protein associations, integrated and transferred across organisms. Nucleic acids research 33, D433–437. 10.1093/nar/gki005.

